# ExSTED microscopy reveals contrasting functions of dopamine and somatostatin CSF-c neurons along the central canal

**DOI:** 10.1101/2021.08.17.456595

**Authors:** Elham Jalalvand, Jonatan Alvelid, Giovanna Coceano, Steven Edwards, Brita Robertson, Sten Grillner, Ilaria Testa

## Abstract

The spatial location of cerebrospinal fluid contacting (CSF-c) neurons enables important regulatory homeostatic functions regarding pH and motion control. Their intricate organization, facing the central canal and extending across the spinal cord, in relation to specific subtypes is poorly understood. This calls for imaging methods with a high spatial resolution (5-10 nm) to resolve the synaptic and ciliary compartments of each individual cell to elucidate their signalling pathways and enough throughput to dissect the cellular organization. Here, light-sheet and expansion microscopy resolved the persistent ventral and lateral organization of dopamine and somatostatin CSF-c neuronal types.

The number of somatostatin-containing dense core vesicles, resolved by STED microscopy, was shown to be markedly reduced upon each exposure to alkaline or acidic pH inhibiting any movement as part of a homeostatic response. Their cilia symmetry was unravelled by ExSTED as sensory in contrast with the motile one found in the dopaminergic ph insensitive neurons. This novel experimental workflow elucidates the functional role of CSF-c neuron subtypes *in situ* paving the way for further spatial and functional cell type classification.

## Introduction

The vertebrate spinal cord contains neurons with many different functions. In the middle there is the central canal containing cerebrospinal fluid. The wall of the central canal is lined with ciliated cells in all vertebrates [1, 2], of which many are GABAergic sometimes with co-transmitters as somatostatin, neurotensin or dopamine [3–6]. Cerebrospinal fluid contacting (CSF-c) neurons [2] located on the lateral parts of the central canal wall express somatostatin and GABA and send axonal processes to mechanosensitive edge cells on the lateral margin of the spinal cord. Cells in a more ventral location contain dopamine [7, 8]. GABA/somatostatin-expressing CSF-c neurons have recently been demonstrated to act as pH and mechanosensors and be part of a pH homeostatic system. At any deviation from neutral pH their activity is increased, which in turn leads to a depression of motor activity [9, 10]. The function of the dopaminergic CSF-c cells is elusive. Additionally, a description of the interplay and fine spatial organization of different CSF-c cell types in the spinal cord is still missing.

In this study, we investigate how somatostatin/GABA and dopaminergic CSF-c neurons are organized throughout the spinal cord with a multi-scale imaging approach, using a high-throughput imaging method with sufficient resolution to resolve individual cells and organelles. Light-sheet microscopy is the method of choice for rapid and minimally invasive acquisitions of large portions of tissues, but with a compromised spatial resolution in favour of recording speed and minimal photo-bleaching. To overcome this problem, we combine light-sheet with expansion microscopy (ExLSM) [11], a technique that physically expands tissue samples by a procedure including sample-embedding in a polyacrylamide gel and ends with ~4–5 fold larger transparent samples [12, 13]. The physical expansion of the sample allows imaging of large volumes of spinal cord tissue with single cell resolution and access to fine spatial information. This imaging approach allows quantification of the relative abundance and spatial patterns of CSF-c neuronal subtypes with specific focus on somatostatin- and dopamine-expressing cells along the central canal.

To further understand the physiological role of dopamine and somatostatin expressing CSF-c neurons, we take advantage of stimulated emission depletion (STED) microscopy to profile their neurotransmitter spatial distribution inside CSF-c neurons. STED microscopy [14, 15] reaches a spatial resolution of ~40 nm, which allows us to resolve single synaptic vesicles even when the organelles are densely packed. This approach can identify neurotransmitter-specific release of dense-core synaptic vesicles during basal activity and upon pH stimulation in both somatostatin- and dopamine-expressing CSF-c neurons.

To gain structural information at ~5–10 nm level, an additional spatial resolution increase is needed, which we demonstrate with the application of STED imaging on expanded spinal cord tissues (ExSTED) [16]. ExSTED imaging features an effective lateral spatial resolution of < 10 nm, and allowed us to obtain cell type-specific structural insight, previously accessible only with EM microscopy, on the cilia subtypes in ciliated somatostatin- and dopamine-expressing CSF-c. The cilia symmetry differs between primary (sensory) cilia (9+0 microtubule duplets) and motile cilia (9+2 microtubule duplets) and could be investigated in the context of specific cell types thanks to the spatial and specificity abilities of ExSTED. We could show that dopamine CSF-c neurons have motile cilia that may contribute to the flow of the cerebrospinal fluid, whereas somatostatin CSF-c neurons instead have predominantly sensory cilia conveying pH and mechanosensitivity.

Overall, this study uses recently developed high resolution imaging techniques adapted to profiling the CSF-c neurons in the spinal cord. This provides cell type information down to the molecular level. The experimental design developed in this study can be applied also to other types of tissue.

## Results

### Distribution of somatostatin- and dopamine-expressing CSF-c neurons along the spinal cord

To investigate the spatial distribution of CSF-c neurons around the central canal at the single-cell level with high speed and sufficient spatial resolution, we combined expansion and light-sheet microscopy on fluorescently labelled CSF-c neurons. An experimental design for expanding and handling spinal cord tissue was developed, which includes slicing, fluorescent immunolabelling of somatostatin and dopaminergic CFS-c neurons in 80–100 μm thick spinal cord slices, and the expansion steps with a final expansion factor of about ~4.5 (Fig. 1a, b).

**Figure 1.**
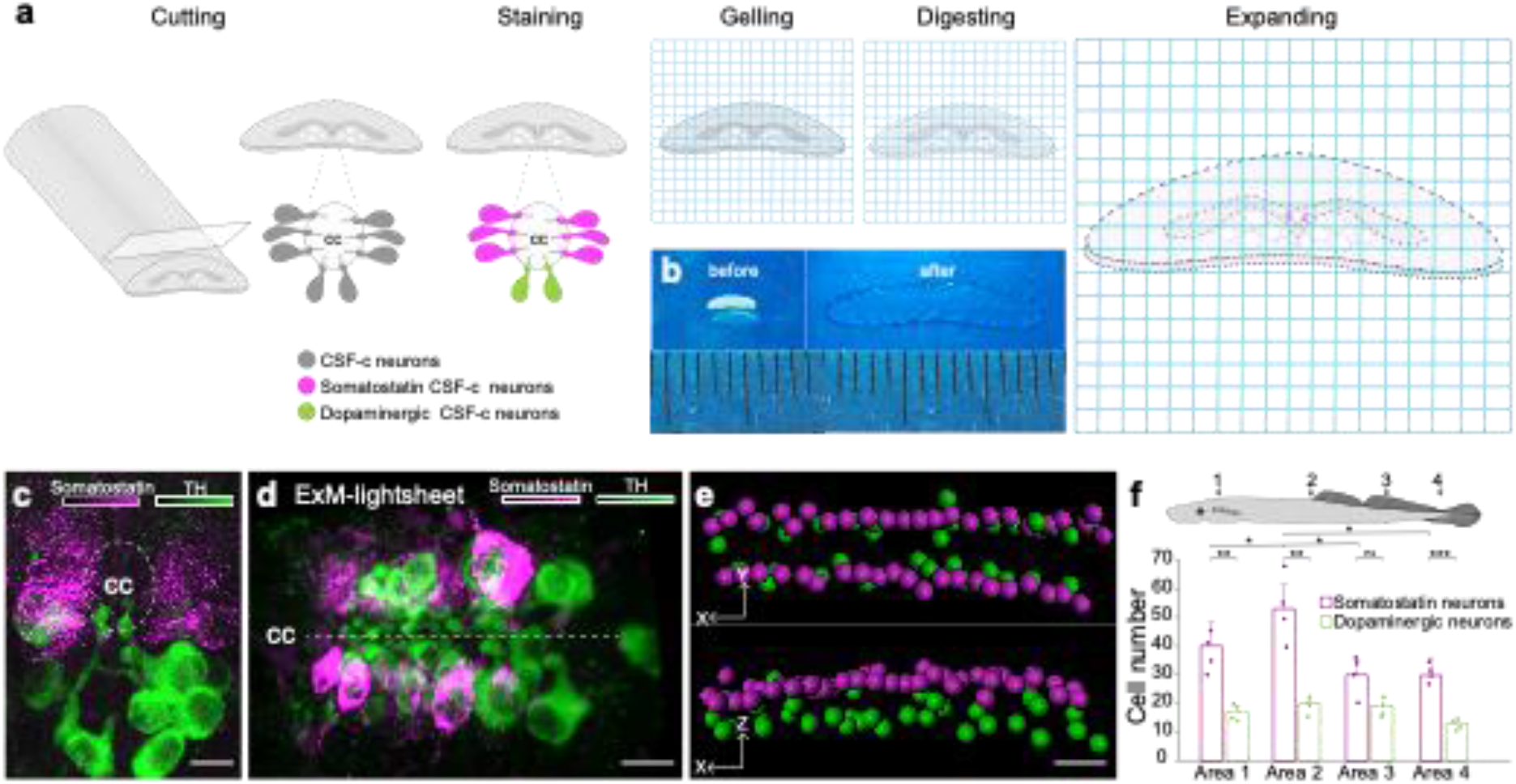
Somatostatin and dopaminergic CSF-c neurons distribution along the spinal cord by expansion and light-sheet microscopy. **a**, A schematic illustration of the lamprey spinal cord treated for expansion microscopy (ExM). The spinal cords were immunostained for somatostatin and tyrosine hydroxylase (TH) prior to the ExM steps (MA-NHS treatment, gelation, proteinase K treatment, and expansion in water). **b**, The spinal cord slices are shown before and after expansion. **c,d**, Expanded samples imaged by light-sheet microscopy along the spinal cord. **c**, Transverse and **d**, horizontal images of somatostatin (magenta) and dopaminergic (green) CSF-c neurons shown by ExM-light-sheet microscopy. Scale bar, 30 μm. **e**, Segmentation of the 3D data from CSF-c neurons. Scale bar, 30 μm. **f**, Quantification of somatostatin- and dopamine-expressing CSF-c neurons in four different areas of the spinal cord. The data are represented as the mean; the error bar represents s.d.; Student’s paired t-test: *p < 0.05 significant difference of somatostatin CSF-c neurons area 1 vs area 2 (*p* = 0.016, *t*_3_ = −4.84), area 2 vs area 3 (*p* = 0.016, *t*_3_ = 5.72) and vs area 4 (*p* = 0.04, *t*_3_ = 3.38), **P < 0.01 and ***P < 0.001 significant difference of somatostatin and dopamine CSF-c neurons at area 1 (*p* = 5.8 × 10^-3^, *t*_3_ = 7.06), at area 2 (*p* = 4 × 10^-3^, *t*_3_ = 7.67), at area 4 (*p* = 7.9 × 10^-4^, *t*_3_ = 13.9), and non-significant difference (n.s.) at area 3 (*p* = 0.09, *t*_3_ = 2.40). cc, central canal.

The combination of our spinal cord expansion protocol and light-sheet microscopy (ExLSM) enables us to record a volume of 360 × 250 × 200 μm^3^ containing a large population of CSF-c neurons where the individual cells can be recognized and counted (Fig. 1c, d). Somatostatin and dopaminergic (TH-expressing) CSF-c neurons can be visualized along the spinal cord (Fig. 1c, d) and their specific location in the 3D architecture of the tissue can be visualized (Supplementary Movie 1) and quantified in different views (Fig. 1e). The cell bodies of both dopamine and somatostatin CSF-c neurons are clearly visible, as well as their characteristic protrusion into the central canal. Different types of CSF-c neurons show a specific distribution along the lamprey spinal cord in all three spatial dimensions. Somatostatin-expressing CSF-c neurons are found throughout the whole volume and located laterally relative to the central canal, while the dopaminergic CSF-c neurons are located ventrally, confirming that the observation on individual sections is maintained in large volume with a high degree of order.

The throughput of ExLSM allowed recording a total volume of 7.2 × 10^7^ μm^3^ along the spinal cord for a total of 224 cells. We quantified the distribution of CSF-c neuronal subtypes at different levels of the spinal cord, resulting in a higher amount of somatostatin CSF-c neurons compared to the dopaminergic CSF-c neurons in all the four areas of the spinal cord. Additionally, in a specific position (area 2) the somatostatin CSF-c neurons were more abundant than in other parts (Fig. 1f).

### Somatostatin and dopamine neurotransmitters in CSF-c neurons are stored in dense core vesicles (DCVs)

To investigate the subcellular location and compartmentalization of somatostatin and dopamine in CSF-c neurons, we used STED microscopy. The STED microscope was equipped with a glycerol objective and red-shifted wavelengths to allow tissue imaging with minimal spherical aberration and scattering.

Somatostatin and dopamine are stored in dense-core vesicles (DCVs) and were found in the soma, bulb protrusion (Fig. 2a–c, e–g), and projections (data not shown) of the CSF-c neurons. No major differences between the somatostatin- and dopamine-positive puncta are observed. In both cases the average diameter of somatostatin and dopamine DCVs was 100–120 nm with vesicles as small as 60 nm (FWHM; Fig. 2d, h) measured with STED. However, when using confocal microscopy, it resulted in an overestimation of vesicle size with 250–300 nm (FWHM; Fig. 2d, h). Our data on the somatostatin and dopamine DCVs diameter recorded with STED are compatible with previously published electron microscopy data [8].

**Figure 2.**
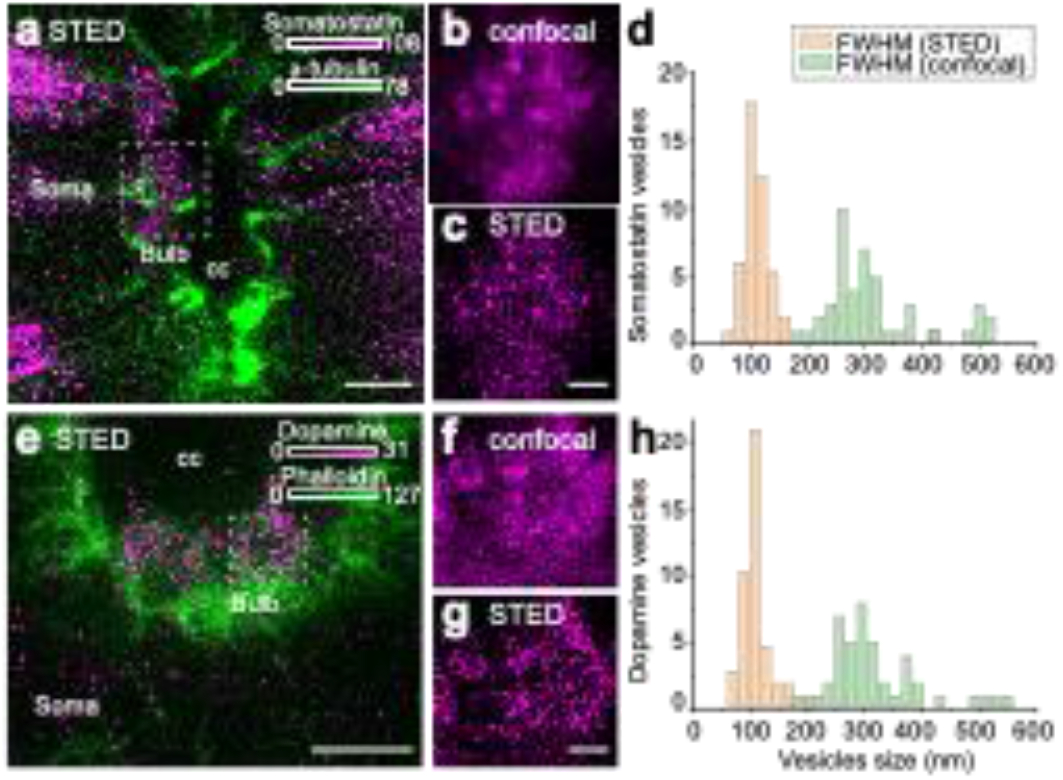
Somatostatin and dopamine in CSF-c neurons are stored in dense core vesicles. **a-c**, Somatostatin (magenta) and α-tubulin (green) immunostaining in CSF-c neurons. Scale bar in a, 5 μm. **b,c**, Selected ROIs of somatostatin DCVs in the bulb of somatostatin-expressing CSF-c neurons imaged with confocal and STED microscopy, respectively. Scale bar, 500 nm. **d**, Analysis of the size of somatostatin DCVs measured with confocal and STED microscopy (n = 46). **e-g**, Dopamine (magenta) and phalloidin (green) immunostaining in CSF-c neurons. Scale bar in **e**, 5 μm. **f,g**, Selected ROIs of dopamine DCVs in the bulb of dopamine CSF-c neurons with confocal and STED microscopy, respectively. Scale bar, 500 nm. **h**, Analysis of the size of dopamine DCVs with confocal and STED (n = 44). cc, central canal.

### Somatostatin but not GABA release are responsible for pH response in somatostatin/GABA CSF-c neurons

The somatostatin CSF-c neurons in the lamprey spinal cord also express GABA [3, 5, 7]. We have shown recently that somatostatin/GABA CSF-c neurons are sensitive to pH changes of the cerebrospinal fluid [9, 10]. Here, we investigate how the spatial distribution and abundance of somatostatin and GABA changed during induced pH changes to acidic (6.5) or alkaline (8.5) conditions and if they were co-released. The somatostatin DCVs were visualized in the soma as well as in the axons by both confocal and STED microscopy (Fig 3a–f). Somatostatin DCVs visualized with STED microscopy were counted in neurons in control, acidic, and alkaline pH conditions. The number of somatostatin DCVs puncta decreased markedly in the soma of CSF-c neurons in either acidic or alkaline pH conditions (Fig. 3a-c and g). In contrast, DCVs increased in the axons (Fig. 3d–f). On the other hand, the slices stained with a GABA antibody did not show any changes in their fluorescent intensity at different pH (Fig. 3h–k).

**Figure 3.**
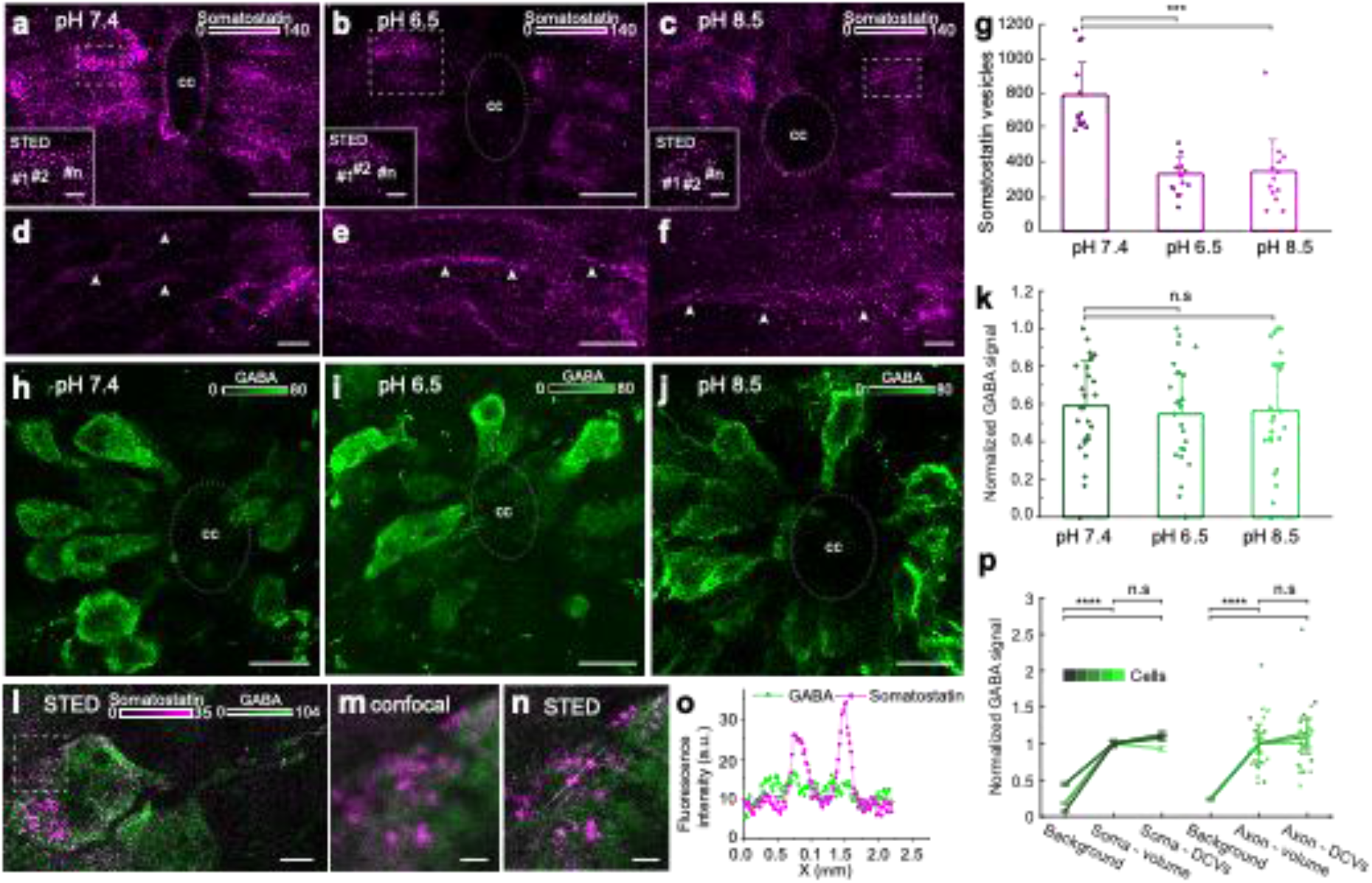
Acidic and alkaline pH decreased the number of somatostatin DCVs in the soma but did not affect GABA intensity. **a-f**, Spinal cord slices in normal (pH 7.4), acidic (pH 6.5), and alkaline (pH 8.5) extracellular solution stained with an anti-somatostatin antibody (magenta). **a-c**, Confocal and STED images (selected ROIs) of somatostatin DCVs in the soma. Scale bar in a-c, 10 μm; in ROIs, 1 μm. **d-f**, The axons of the somatostatin-expressing CSF-c neurons (arrowheads). Scale bar, 10 μm. **g**, Quantification of somatostatin DCVs in the different conditions (n = 13). Student’s paired t-test: ***p < 0.001 significant difference between pH 7.4 and 6.5 (*p* = 1.9 × 10^-6^, *t*_12_ = 8.5), and 7.4 and 8.5 (*p* = 1.7 × 10^-4^, *t*_12_ = 5.3). **h-j**, The spinal cord slices in normal, acidic and alkaline extracellular solution, stained with an anti-GABA antibody (green). Scale bar, 10 μm. **k**, Comparison of normalized GABA signals at pH 7.4 (n = 26), 6.5 (n = 24), and 8.5 (n = 22), respectively. Student’s t-test: non-significant difference (n.s.) between pH 7.4 and 6.5 (*p* = 0.62, *t*_47_ = 0.48), and 7.4 and 8.5 (*p* = 0.80, *t*_43_ = 0.25). **l-o**, STED and confocal images of spinal cord slices stained for somatostatin (magenta) and GABA (green). **l**, STED image of a CSF-c neuron. Scale bar, 1 μm. **m,n**, Selected ROI from the soma in **l** shown at higher magnification with confocal and STED microscopy, respectively. Scale bar, 0.3 μm. **o**, Line profile graph in image n. **p**, Mean GABA signal in cellular compartments and compared to extracellular background (n=5), normalized to volume intensity in soma (n=5) and axons (n=3) respectively. Repetitions are different cells. Student’s t-test between means of cellular means: **** p < 0.0001 significant difference between soma-volume and background (*p* = 1.0 × 10^-5^, *t*_8_ = −9.7), soma-DCVs and background (*p* = 1.0 × 10^-5^, *t*_8_ = −9.6), axon-volume and background (*p* = 8.0 × 10^-8^, *t*_4_ = −93), and axon-DCVs and background (*p* = 4.4 × 10^-5^, *t*_4_ = −19), non-significant differences (n.s.) between soma-volume and DCVs (*p* = 0.090, *t*_8_ = −1.9), and axon-volume and DCVs (*p* = 0.12, *t*_4_ = 1.9). Data (**g, k, p**) is represented as means, with error bars representing s.d. (**g, k**) or s.e.m. (**p**). cc, central canal.

As somatostatin and GABA are co-expressed in the same CSF-c neurons, we investigated if they were colocalized in the same vesicles (Fig. 3l–p). STED images of somatostatin DCVs and GABA-expressing CSF-c neurons did not show colocalization of somatostatin DCVs and GABA in the soma (Fig. 3m, n, and o). GABA signal showed no significant differences when measured in and outside of somatostatin DCVs, neither at the cell body (soma) nor at the axons (Fig. 3p). The results confirmed that there is no correlation between GABA and somatostatin signal in somatostatin vesicles. Our imaging data support the conclusion that during pH changes of the extracellular solution vesicles containing somatostatin are released, resulting in fewer DCVs in the cell bodies, while GABA is not coreleased.

### Dopaminergic CSF-c neurons are not sensitive to changes in extracellular pH

The next step was to explore whether the dopaminergic CSF-c neurons at the ventral part of the central canal were also sensitive to pH changes. As for the somatostatin experiments, acidic and alkaline extracellular pH was perfused on spinal cord slices and subsequently stained with a dopamine antibody to investigate the dopamine DCVs spatial distribution in dopaminergic CSF-c neurons. We performed STED microscopy to resolve and count the single vesicles in distinct neuronal locations (Fig. 4a–c). The result showed no significant change in the number of dopaminergic DCVs after perfusion with acidic or alkaline pH (Fig. 4d).

**Figure 4.**
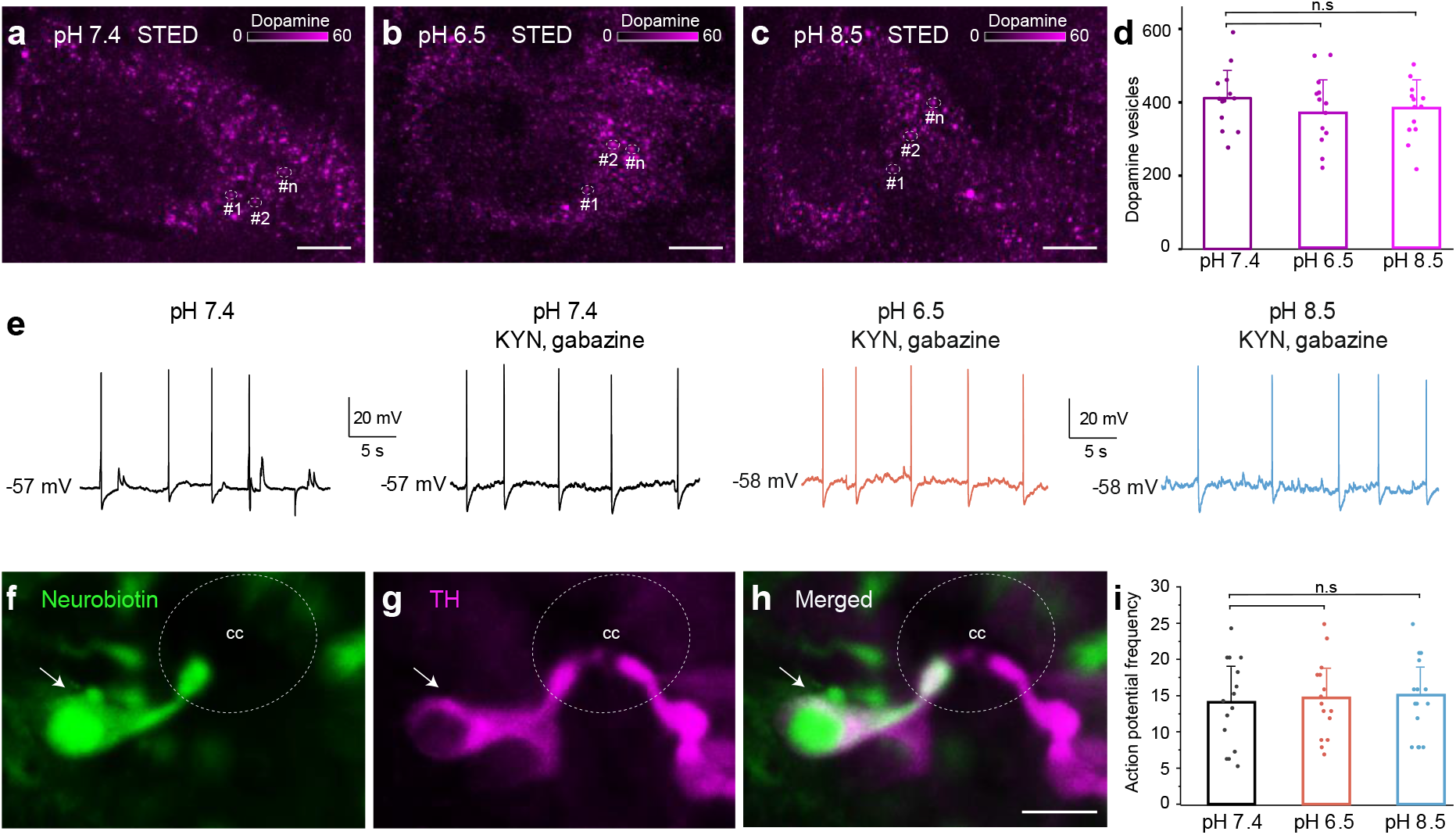
Dopaminergic CSF-c neurons did not respond to acidic and alkaline pH. **a-c**, STED images of dopamine-containing DCVs in the soma of CSF-c neurons in normal (pH 7.4), acidic (pH 6.5), and alkaline (pH 8.5) extracellular solution. Scale bar 1 μm. **d**, Quantification of the number of dopamine DCVs in the soma of CSF-c neurons in the different conditions at the same total cells (n = 13). Student’s t-test: non-significant (n.s.) between pH 7.4 and 6.5 (*p* = 0.33, *t*_12_ = 1), and 7.4 and 8.5 (*p* = 0.62, *t*_12_ = 0.50). **e**, Whole-cell patch recording of a CSF-c neuron, showing firing spontaneous action potentials in control (pH 7.4), acidic (p H 6.5) and alkaline (pH 8.5) conditions in the presence of gabazine (20 mM) and kynurenic acid (2 mM). **f-h**, Photomicrographs of the CSF-c neurons recorded in **e** intracellularly filled with Neurobiotin (arrow) during recording. The labelled cell showed immunoreactivity to TH (arrow). Scale bar, 10 μm. **i**, Action potential frequency during 1 min in CSF-c neurons at pH 7.4, 6.5, and 6.8, respectively (n = 15). Student’s paired t-test: non-significant difference (n.s.) between pH 7.4 and 6.5 (*p* = 0.24, *t*_14_ = −1.22), and 7.4 and 8.5 (*p* = 0.1, *t*_14_ = −1.75). The bar graph data are represented as the means, with error bars representing s.d. cc, central canal.

To complement the results from the STED imaging, dopaminergic CSF-c neurons were patched as previously described for somatostatin-expressing CSF-c neurons [9, 10]. Their electrophysiological properties and response to changes of extracellular pH were investigated with whole-cell patch recording in current-clamp mode (Fig. 4e). All recorded dopaminergic CSF-c neurons fired spontaneous action potentials, depolarizing synaptic potentials, and hyperpolarizing synaptic potentials (Fig. 4e). To verify that responses were not evoked synaptically, the GABAergic and glutamatergic synaptic transmission was blocked by bath-application of gabazine (GABA_A_ receptor antagonist) and kynurenic acid (glutamate receptor antagonist). During bath-applied extracellular solutions of acidic pH (6.5) or alkaline pH (8.5) no changes in spike frequency was observed, nor a net depolarization of the resting membrane potential (Fig. 4e and i). Thus, in contrast to the somatostatin CSF-c neurons the dopamine CSF-c are not sensitive to pH changes. All neurons labelled with Neurobiotin during the patch-clamp recording were TH-immunoreactive (Fig. 4f–h).

Both somatostatin and dopamine CSF-c neurons are ciliated (see below) and the former known to be mechanosensitive [10]. To test if the dopaminergic CSF-c neurons are mechanosensitive very brief fluid pulses were applied near their bulb protrusion in the central canal as previously done for the somatostatin CSF-c neurons [10]. Dopamine CSF-c neurons responded with an action potential and a distinct mechanosensitive response (Extended Data Fig. 1a). To investigate if the mechanosensitivity in dopaminergic CSF-c neuron was mediated by the acid-sensing ion channel 3 (ASIC3) as in somatostatin CSF-c neurons, APETx2, a specific blocker of ASIC3, was applied. In the presence of APETx2 we still observed a mechanosensitive response (Extended Data Fig. 1b). Thus, the mechanosensitivity is not mediated by ASIC3 in dopamine CSF-c neurons. *In situ* hybridisation showed expression of Polycystic kidney disease 2-like 1 (PKD2L1) channels in dopaminergic CSF-c neurons (Extended Data Fig. 1c). As PKD2L1 has been confirmed as a mechanosensitive ion channel in zebrafish [17], the results suggest that the mechanosensitivity of dopaminergic CSF-c neurons may be mediated by PKD2L1 channels.

In conclusion, dopamine CSF-c neurons are not sensing pH changes as their somatostatin counterparts, but both are mechanosensitive while the transduction is mediated by ASIC3 in the latter and by PKD2L1 in the dopamine CSF-c neurons.

### CSF-c neurons show both primary and motile cilia symmetries

Somatostatin- and dopamine-expressing CSF-c neurons are thus both mechanosensitive, but through different transduction mechanisms. Since both types of CSF-c neurons are ciliated, we investigated whether the functional difference is reflected in the structural organization of the cilia.

Cilia express acetylated α-tubulin and can be classified either as primary cilia, mainly present in sensory cells and neurons and with a 9+0 symmetry of nine outer microtubule doublets, or motile cilia, with a 9+2 symmetry showing an extra pair of microtubule singlets in the centre [18, 19]. Primary and motile cilia both have an average diameter of 200–240 nm, close to the achievable spatial resolution of a confocal microscope equipped with a high numerical aperture objective. Therefore, a higher spatial resolution is crucial to assess their organization and sub-compartmentalisation.

One aim for exploring the cilia symmetry of ciliated CSF-c neurons was to uncover potential signalling compartments related to pH and mechanosensitivity of the CSF-c neurons. Cilia in the central canal was immunolabelled for acetylated α-tubulin, a protein characteristic of cilia, and imaged with confocal and STED microscopy to measure their diameters, respectively (Fig. 5a, b, e, and f). STED imaging visualises the cilium as a hollow structure with an outer diameter of about 240 nm thanks to the increased spatial resolution, but the resolution is still not enough to detect the microtubule doublets. One strategy to increase the observable level of detail is to expand the tissue. In the expanded spinal cord tissue, pre-immunostained for acetylated α-tubulin, the resolution of the dense cilia in the central canal increased (Extended Data Fig. 2, 3D XYZ image) as compared to confocal imaging of the non-expanded sample. Confocal imaging of the expanded sample shows spatial details comparable to STED images (Fig. 5c) of the non-expanded one, therefore still not having the resolution (5–10 nm) needed to dissect the internal cilia structure and identify microtubule doublets. That level of detail is crucial to further investigate the nature of cilia as sensory or motile for each specific cell type.

**Figure 5.**
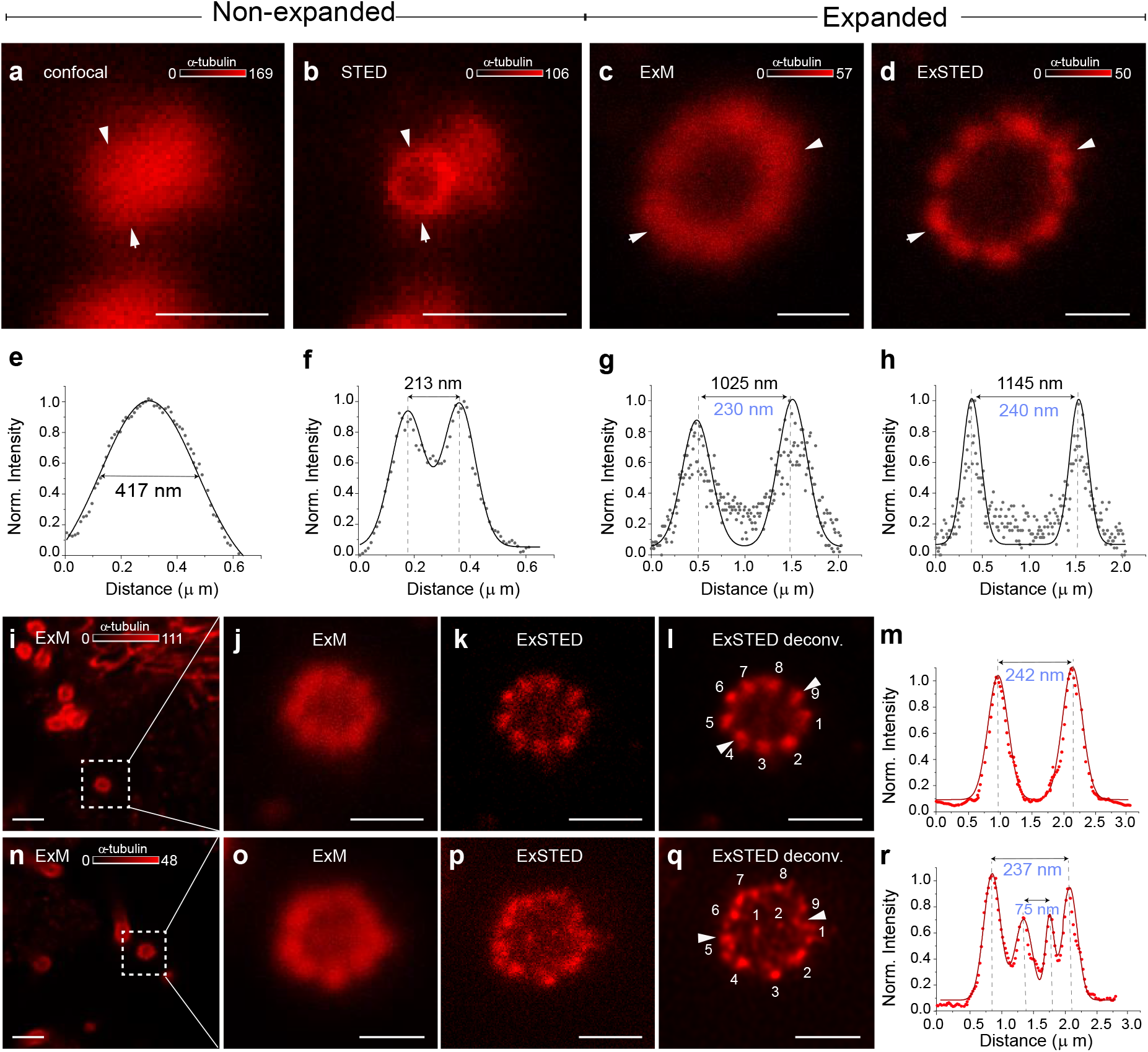
Primary and motile cilia symmetry are present in the lamprey spinal cord. **a-b**, Confocal and STED images of a CSF-c neuron cilium in a non-expanded spinal cord pre-stained with anti-α-tubulin antibodies. Scale bar, 0.5 μm. **c,d**, Confocal (ExM) and STED (ExSTED) images of a cilium in the expanded spinal cord. Scale bar, 0.5 μm. **e,f**, Quantification of the cilium diameter (arrowheads) in confocal and STED images in a non-expanded spinal cord. **g,h**, Quantification of the cilium diameter (arrowheads) in confocal (ExM) and STED (ExSTED) in the expanded spinal cord. **i-m**, Confocal (ExM, **i**, **j**) and STED (ExSTED, **k,l** deconvoluted) images of a primary cilium with 9+0 symmetry in the expanded spinal cord. Scale bar i, 2 μm, j-l, 1 μm. **m**, Quantification of the cilium diameter from image **l** (arrowheads). **n-r**, Confocal (ExM, **n,o**) and STED (ExSTED, **p,q**, deconvoluted) images of a motile cilium with 9+2 symmetry in the expanded spinal cord. Scale bar n, 2 μm, o-q, 1 μm. **r**, Quantification of the cilium diameter from image **q** (arrowheads). Cilia diameters in blue were divided by the expansion factor.

We then used the combination of STED microscopy and expanded tissue (ExSTED). With ExSTED we were able to add a factor of ~4–5 to the typical 50 nm resolution of STED microscopy and therefore resolving at the smaller spatial scale of 5–10 nm. In this way it was possible to separate high-density cilia within the 3D geometry of the central canal (Supplementary Movie 2) due to the increased imaging resolution and further resolve their internal structure, allowing to classify them as motile or sensory (Fig. 5d). In some cilia, the central pair of tubules was clearly observed with a peak-to-peak distance of 70 nm (Fig. 5p-r).

We explored both types of cilia symmetries in the lamprey spinal cord: sensory (Fig. 5i-m) and motile cilia symmetries (Fig. 5n-r). The presence of motile cilia in the central canal of the lamprey has been shown with electron microscopy [2, 8], but with ExSTED we could detect both motile and sensory cilia symmetries in this area. As all CSF-c neurons, including somatostatin and dopaminergic CSF-c neurons in the lamprey spinal cord, are ciliated, the next step is to know which cilia type is related to which neuronal phenotypes and if they contribute to a specific neuronal function.

### Somatostatin CSF-c neurons have both primary and motile cilia, but dopaminergic CSF-c neurons have only motile cilia

To further investigate whether somatostatin-expressing CSF-c neurons (pH-sensitive) and dopaminergic CSF-c neurons (non-pH-sensitive) have different types of cilia, expansion was combined with STED microscopy. Using dual-colour imaging of spinal cord slices, either somatostatin or dopamine (TH immunostaining), together with α-tubulin immunostaining (Fig. 6) was analysed. To have better access to the cilia and CSF-c neurons the central canal of the expanded spinal cord has been cut horizontally and flipped 90° with respect to the coverslip. The central canal can then be visualized in a cylindrical form (Figure 6a–c). To confirm what type of CSF-c neurons the cilia belong to, z-stack scanning for all the imaging was applied. In our data, 70 cilia belong to somatostatin-expressing CSF-c neurons (n=70). Of these, 60 cilia were primary (9+0 symmetry) and 10 cilia were motile (9+2 symmetry) (Fig. 6d–k and t). Additionally, 20 cilia of dopaminergic CSF-c neurons were all motile cilia (9+2 symmetry) (Fig. 6l-s and t). We found that somatostatin-expressing CSF-c neurons are sensory neurons expressing mainly sensory cilia. However, the motile cilia in both somatostatin-expressing and dopaminergic CSF-c neurons might be involved in contributing to the flow of CSF through the central canal (Fig. 6u).

**Figure 6.**
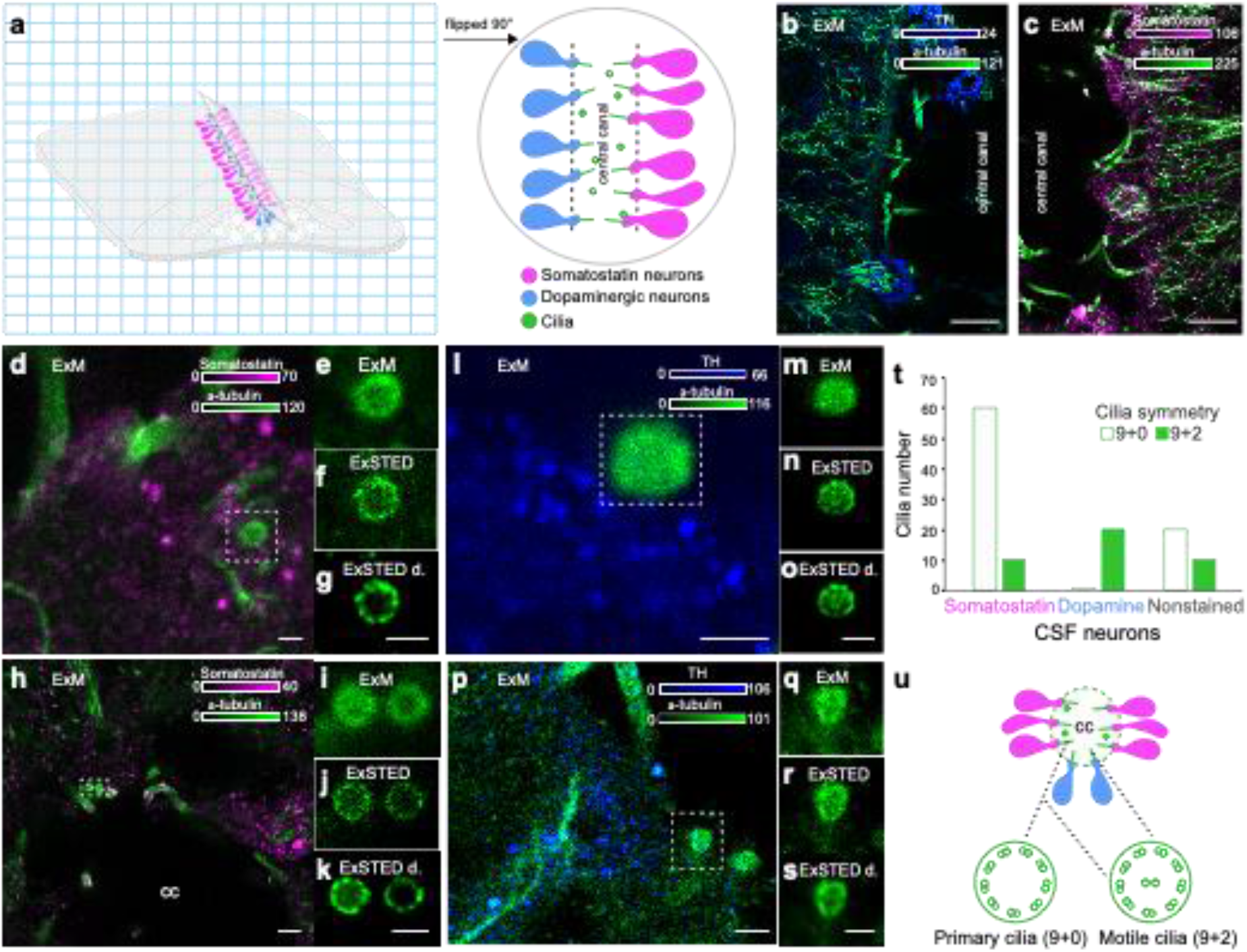
Cilia symmetries in somatostatin and dopaminergic CSF-c neurons. **a**, A schematic illustration of an expanded spinal cord stained for somatostatin (magenta), dopamine (blue), and α-tubulin (green). The gel was cut through the central canal and flipped 90° on the side on the coverslip. **b,c**, Longitudinal images of the expanded spinal cord (ExM) stained with α-tubulin, TH, and somatostatin antibodies, respectively. Scale bar, 10 μm. **d-k**, ExM and ExSTED images of two somatostatin CSF-c neurons with their cilia. Scale bar d, 1 μm, h, 3 μm. **f,g**, showing 9+0 symmetry and **j,k**, showing both 9+0 and 9+2 symmetries. Scale bar, 1 μm. **l-s**, ExM and ExSTED images of two dopaminergic CSF-c neurons (TH staining) with 9+2 symmetry. Scale bar, l-o, q-s, 1 μm, p, 2 μm. **t**, Quantification of cilium types in somatostatin and dopaminergic CSF-c neurons. **u**, A schematic illustration of the central canal with somatostatin and dopaminergic CSF-c neurons and their possible cilia symmetries. cc, central canal.

The sensory cilia are a potential location for pH-sensitive receptors [20, 21]. We have recently shown that in somatostatin-expressing CSF-c neurons, acidic and alkaline responses are mediated by the acid-sensing ion channel 3 (ASIC3) and polycystic kidney disease 2-like 1 (PKD2L1) channel, respectively [9, 10]. Interestingly, we could detect the expression of both ASIC3 and PKD2L1 on the cilia of spinal CSF-c neurons in mice (Extended Data Fig. 3a-q). Besides, we visualized Arl13b, a ciliary protein on cilia that has high expression in sensory cilia (Extended Data Fig. 3r-v).

### CSF neurons have one and rarely two cilia

Another advantage of expansion microscopy (ExM) is that more specific morphological details of cells can be revealed, which cannot be detected in non-expanded samples. Using ExM, paired cilia of CSF-c neurons in their bulb protrusion into the central canal were visualised in some cases (Fig. 7). As reported, somatostatin-expressing and dopaminergic CSF-c neurons mainly have one cilium, but we could in rare cases (N=8) detect two cilia per bulb of these neurons by ExM (Fig. 7a-f). To confirm the finding, we performed z-stack scanning to see the whole bulb and its cilia in 3D (Supplementary Movie 3).

**Figure 7.**
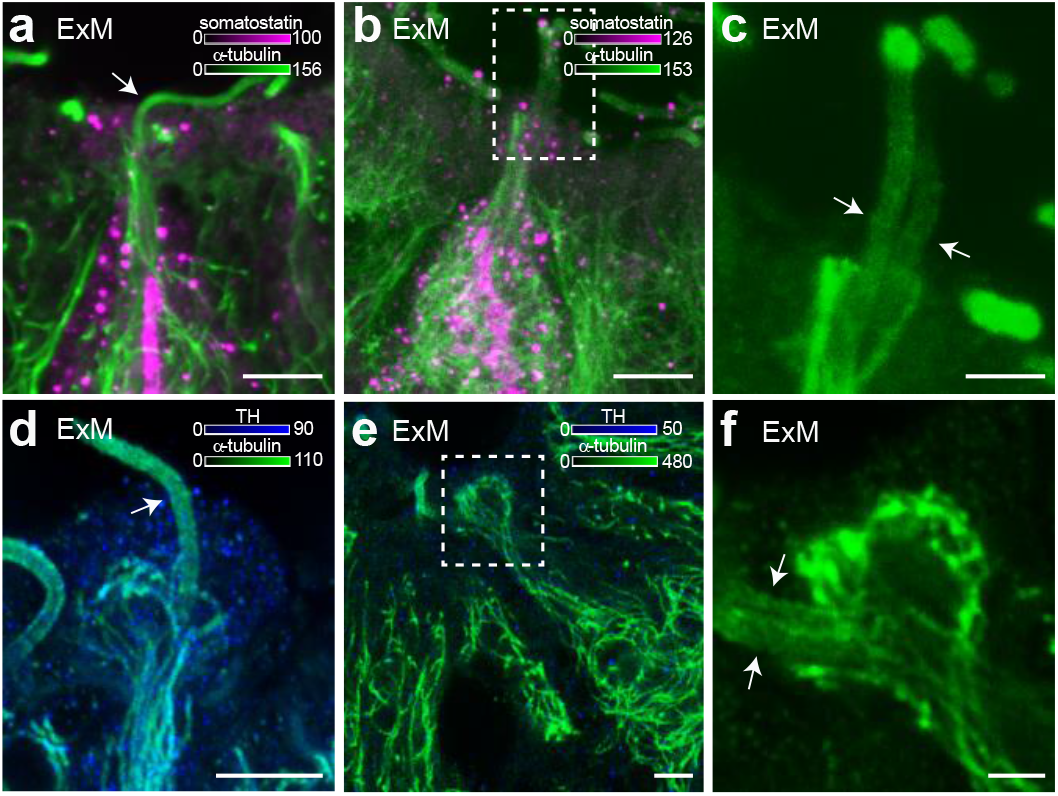
CSF-c neurons have one or two cilia on their bulb protrusions. **a**, An ExM image of an expanded somatostatin CSF-c neuron showing one cilium on its bulb (arrow). Scale bar, 5 μm. **b,c**, A somatostatin CSF-c neuron with two cilia (arrow) (**c** selected ROI from **b**). Scale bar b, 5μm and c, 2μm. **d**, A dopaminergic CSF-c neuron (TH expressing) with one cilium (arrow). Scale bar, 5 μm. **e,f**, A dopaminergic CSF-c neuron with two cilia (arrows) (**f** selected ROI from **e**). Scale bar e, 5μm and f, 2μm. cc, central canal.

## Discussion

In this study, we uncovered the dense spatial organization of somatostatin- and dopamine-expressing CSF-c neurons along the central canal. ExLSM allowed us to visualize and quantify CSF-c neurons in a large volume of the spinal cord with a fourfold increase in resolution, necessary to distinguish and count each individual cell. There was approximately twice as many somatostatin CSF-c neurons as dopaminergic CSF-c neurons along the whole length of the spinal cord. One area closest to rostral dorsal fin had, however, a somewhat larger number of somatostatin CSF-c neurons than elsewhere. This is the same spinal cord area in which a dramatic decrease occurs of the serotonin/dopamine neurons that are located just ventral to the central canal [22, 23]. Whether this is a coincidence or a related phenomenon remains to be elucidated.

We were able to quantify somatostatin and dopamine DCVs with STED microscopy. The size distribution is in line with previous electron microscopy studies [5, 8]. The STED data reveal the presence of DCVs in several CSF-c cellular sub-compartments, including the bulb protrusion into the central canal, soma, and axonal branches. We could show a marked reduction of somatostatin DCVs in the soma of CSF-c neurons when exposed to acidic or alkaline pH, while there was no effect on the number of dopamine DCVs. We have shown previously that somatostatin/GABA-expressing CSF-c neurons are responding to acidic and alkaline extracellular solutions [9, 10] being part of a homeostatic system for reducing motor activity when the spinal cord/brain is exposed to changes of pH as due to metabolic or respiratory stress or intense muscle activity. Interestingly, the STED imaging experiments showed that the pH response resulted from somatostatin DCVs release, whereas GABA was unaffected. The dopaminergic CSF-c neurons, in contrast to somatostatin-expressing CSF-c neurons, were not influenced by changes of pH.

At the ultrastructural level, GABAergic CSF-c neurons have been shown to have ciliated endings in the lamprey and other species [2, 8, 24]. We show here that somatostatin/GABA and dopaminergic/GABA CSF-c neurons have similar responses to mechanical stimuli, but only the former responds to pH changes. The question arose if these two types of CSF-c neurons have different forms of cilia.

To visualize cilia microtubule symmetry in CSF-c neurons a resolution of 5–10 nm is needed, which was provided by the combination of expansion and STED microscopy (ExSTED) [16]. ExSTED microscopy allowed us to obtain cell type-specific structural insights on the cilium types in ciliated somatostatin- and dopamine-expressing CSF-c neurons, respectively. We found both primary cilia with 9+0 symmetry, which may sense a wide variety of signals and are found in specialized sensory cells [25, 26], and motile cilia with 9+2 symmetry, which mainly generate fluid flow [27, 28].

A high expression of sensory cilia (9+0) was found in pH/mechanosensitive somatostatin-expressing CSF-c neurons, which provides an additional insight to recognize these neurons as sensory neurons. The ion channel (ASIC3) mediates both the acid-sensing and the mechanosensitivity, and PKD2L1 the alkaline response in somatostatin-expressing CSF-c neurons [9, 10]. We have also observed expression of ASIC3 and PKD2L1 channels as well as sensory cilia in somatostatin CSF-c neurons of the mouse, which suggests that this may be representative of all vertebrates. On the other hand, the motile cilia (9+2) were found in the non-pH-but mechanosensitive dopaminergic CSF-c neurons, which may possibly suggest that these neurons act as a CSF flow generator. However, the expression of the PKD2L1 channel as a mechanosensitive receptor [29, 30] in dopaminergic CSF-c neurons might be involved in their mechanosensitive response. The mechanosensitive of GABA CSF-c neurons sense the lateral bending movements occurring during locomotion [9, 31] and has also been shown to modulate locomotor movements in the zebrafish.

In conclusion, by applying expansion microscopy combined with light-sheet and STED microscopy with nanoscale precision, the spatial organization, abundance, and sub-cellular composition of two distinct GABAergic CSF-c neuronal subtypes in the lamprey spinal cord were elucidated, leading to a deeper understanding of their physiological role.

## Online Material and methods

### Animals

#### Lamprey

Experiments were performed on a total of 40 adult river lampreys (*Lampetra fluviatilis*) of both sexes that were collected from the Ljusnan River, Hälsingland, Sweden. The experimental procedures were approved by the local ethical committee (Stockholms Norra Djurförsöksetiska Nämnd) and were in accordance with The Guide for the Care and Use of Laboratory Animals (National Institutes of Health, 1996 revision). During the investigation, every effort was made to minimize animal suffering and to reduce the number of animals used during the study.

#### Mouse

Experiments were performed on a total of 4, c57 wild type mice. All experiments were performed in accordance with animal welfare guidelines set forth by Karolinska Institutet and were approved by Stockholm North Ethical Evaluation Board for Animal Research.

### Immunohistochemistry

#### Lamprey

The animals (n=30) were deeply anesthetised through immersion in carbonate-buffered tap water containing MS-222 (100 mg/L; Sigma, St Louis, MO, USA). Following decapitation, a portion of the spinal cord, rostral to the dorsal fin, was fixed by 4% (w/v) paraformaldehyde (PFA) in PBS for 12–24 h at 4°C, and subsequently cryoprotected in 20% sucrose in phosphate buffer (PB) for 3–12 h. For GABA and dopamine immunodetection 1% glutaraldehyde (v/v) was added to the fixative solution. Transverse sections (20 μm thick) were cut on a cryostat (Microm HM 560, Microm International GmbH, Walldorf, Germany), collected on gelatine-coated slides, and kept at −20°C until processing.

Sections were incubated/co-incubated with different primary antibodies listed here: a mouse monoclonal anti-acetylated tubulin antibody (dilution 1:500, Sigma-Aldrich), a rat monoclonal anti-somatostatin antibody (1:100, Millipore) a rabbit polyclonal anti-somatostatin-14 IgG antibody (1:1000, Peninsula laboratories international), a mouse monoclonal anti-TH antibody (1:200, Millipore), a rabbit polyclonal anti-tyrosine hydroxylase (TH) antibody (1:500, Sigma-Aldrich), a mouse monoclonal anti-dopamine antibody (1:400, Sigma-Aldrich) and/or a mouse monoclonal anti-GABA antibody (1:2000, Swant).

#### Mouse

The animals were deeply anesthetised with sodium pentobarbital (200 mg/kg i.p.) and transcardially perfused with 4% PFA in 0.01 M phosphate buffered saline (PBS) pH 7.4 The spinal cord was removed and postfixed for 2 hours, after which it was transferred to a 12% sucrose solution in 0.01M PBS overnight. Transverse sections (20 μm thick) were cut on a cryostat and mounted on gelatine-coated slides and kept at −20°C until processing. Sections were incubated with a rabbit polyclonal anti-polycystin-L antibody (1:500, Merk Millipore), a rabbit polyclonal ASIC3 antibody (1:400, TermoFisher Scientific), or a rabbit polyclonal anti-ARL13B antibody (1:500, Proteintech).

The lamprey and mouse spinal cord sections were after incubation with primary antibodies thoroughly rinsed in PBS and then incubated with Alexa Fluor 594-conjugated donkey anti-rat IgG (1:500, Jackson ImmunoResearch) or Alexa Fluor 488-conjugated donkey anti-rat IgG (1:200, Jackson ImmunoResearch), STAR 635P-conjugated goat anti-mouse (1:500, Abberior) or Alexa Fluor 594-conjugated donkey anti-mouse (1:500, TermoFisher Scientific), and STAR 635P-conjugated goat anti-rabbit (1:500, Abberior) or Alexa Fluor 594-conjugated goat anti-rabbit (1:500, TermoFisher Scientific), for 3 hours at room temperature. The sections were Nissl stained by adding NeuroTrace^™^ 530/615 red or 640/660 deep-red fluorescent Nissl stain (1:1000, Invitrogen) to the secondary antibody solution. Phalloidin conjugated with STAR 635P (1:200, Abberior) was added to the secondary antibodies to stain actin filaments. The primary and secondary antibodies were diluted in 1% bovine serum albumin, 0.3% Triton X-100 in 0.1 M PB. All sections were mounted in custom-made Mowiol mounting media, supplemented with DABCO (Thomas Scientific, C966M75), and covered with coverslips (No. 1.5).

### Expansion microscopy

After fixation, transverse sections of the lamprey spinal cord (40–50 μm thick) were cut on a cryostat and collected and immersed in PBS. The sections were stained with antibodies according to the protocols described above. The samples were treated at room temperature for 1 hour with anchoring solution, 1 mM MA-NHS in PBS (0.018 g MA-NHS in 100 μl DMSO and stored at −20°C) which enables proteins to be anchored to the hydrogel. The slices then were incubated for 1 hour in a monomer solution on ice, which was followed by adding the gelling solution to the gelling chambers. The gelling solution consisted of monomer solution (1 M NaCl, 8.6% sodium acrylate, 2.5% acrylamide and 0.15% N,N’-methylene bisacrylamide in PBS), 4-hydroxy-TEMPO (0.2%), TEMED (accelerator solution, 0.2%) and APS (initiator solution, 0.2%). The slices were incubated in a 37°C incubator for 2 hours for gelation. The gels were removed from the gelling chamber and the coverslips were transferred to digesting buffer (50 mM Tris pH 8.0, 1mM EDTA, and 0.5 Triton X-100) with proteinase K (1:100, 8 units/ml, New England Biolabs) for 30–40 min in room temperature. The gels were removed from the digestion buffer and immersed in deionized (DI) water (3–5 times for 30 min) for further expansion. After final expansion, the gels were mounted to a 35 mm glass bottom petri dish coated with poly-L-lysine (Sigma-Aldrich). To remove the extra gel on a coverslip and increase the resolution, we cut the gel through the central canal and rotated it 90 degrees away from the coverslip. The expansion factor was ^~^4.5–5 and has been calculated by overlapping the pre- and post-expanded gel slice in the air-water boundary.

### STED microscopy

Most STED images have been recorded on a custom-built setup, as previously described [32] equipped with a glycerol objective (HCX PL APO 93x/1.30 NA GLYC STED White motCORR, 15506417, Leica Microsystems). The images were recorded by exciting Alexa Fluor 594 and STAR 635P with 561 nm (PDL561, Abberior Instruments) and 640 nm (LDH-D-C-640, PicoQuant) laser lines, respectively. A STED beam at 775 nm (KATANA 08 HP, OneFive) has been used to deplete both laser lines, shaped by a spatial light modulator (LCOS-SLM X10468-02, Hamamatsu Photonics) into a donut. Two-colour STED images were recorded line-by-line sequentially. A third confocal channel, for structures labelled with Alexa Fluor 488, has been excited with a 510 nm laser line (LDH-D-C-510, PicoQuant). Detection was performed using the following bandpass filters: ET615/30m (Chroma), ET705/100m (Chroma), FF01-550/49 (Semrock). The pixel sizes of the images were 27.2-32.3 nm in XY, and 300 nm in Z. The pixel dwell time used was 10–50 μs, added over 1–2 lines. Laser powers used were in the following ranges: 561 nm: 1–40 μW, 640 nm: 2–20 μW, 775 nm: 65–180 mW. Additional STED images (Figure 2 and Figure 3a–j) have been recorded on a commercial Leica TCS SP8 3X STED. The images were recorded by exciting Alexa Fluor 488, Alexa Fluor 594, and STAR 635P with laser lines at 488 nm, 594 nm, and 635 nm, respectively. The detection windows used were 510–560 nm, 600–645 nm, and 670–750 nm. The pixel size of the images was 25–30 nm, and the pixel dwell time used was 10–40 μs, added over 2–4 lines.

### Light-sheet microscopy

The expanded slices from four different parts of the lamprey spinal cord were separately glued to a metal rod and placed in the chamber of a Zeiss Light-sheet Z1 microscope containing DI water. Fluorescence was excited from two sides using 10x/0.2 NA illumination objectives and detected using a 10x/0.4 NA or 20x/1.0 NA water dipping objective.

### Image analysis

The images were analyzed using Fiji [33] and Imspector (Max-Planck Innovation) and MATLAB (The Mathworks). The transverse sections of the cilia images were deconvoluted with a calculated effective STED PSF, Lorentzian with FWHM of 50 nm, using the Richardson-Lucy deconvolution implemented in Imspector. The deconvolution was stopped after 5 iterations. The line profiles of cilia and DCVs were fitted with a Gaussian, and the full width at half maximum (FWHM) was measured. The OriginLab software (OriginLab) was used for making the graphs. Animation and spot object creation tools of Imaris 9.1 (Bitplane) were used to make 3D movies and segmentation of the 3D data.

Analysis of the correlation of GABA signal and somatostatin DCVs (Figure 3v) was performed by manually selecting areas inside (Soma/Axon – DCVs) and outside (Soma/Axon – volume) somatostatin DCVs (detected in the somatostatin channel) as well as in the extracellular space (Background). The mean of the GABA signal inside each area was recorded. A total of 5 cells were used for the soma and 3 cells for the axons. The number of areas selected in each category in each cell ranged from 3 to 63, with a mean of 37. Plotted are the paired graphs for each cell, with each data point representing the cellular mean of the mean GABA signals per area. The Student’s t-test is performed between the groups of means of the cells per category.

### Live slices

The spinal cord of lamprey (n = 5) was dissected out and embedded in 4% agar dissolved in ice-cooled oxygenated HEPES-buffered physiological solution containing (in mM): 138 NaCl, 2.1 KCl, 1.8 CaCl2, 1.2 MgCl2, 4 glucose, 2 HEPES, and with pH adjusted to 7.4 with NaOH). The agar block containing the spinal cord was glued to a metal plate and transverse slices of the spinal cord (100–150 or 300 μm) were cut on a vibrating microtome. The preparation was continuously perfused with HEPES solution at 4–6 °C. Then the spinal cord slice was exposed to HEPES solution with various pH values (7.4, 6.5, 8.5) for 8–10 min. The slices were then fixed immediately with 4% PFA (somatostatin) or 4% PFA and 1% glutaraldehyde (dopamine and GABA) in 0.01 M PBS overnight at 4°C. Following a thorough rinse in 0.01 M PBS, the slices (100–150 μm) were incubated with a rat monoclonal anti-somatostatin, a mouse monoclonal anti-GABA, or a mouse monoclonal anti-dopamine antibodies overnight at 4°C. The slices were then incubated with Alexa Fluor 594-conjugated donkey anti-rat or anti-mouse IgG as described above.

### Patch-clamp recordings

The spinal cord slices (300 μm) of lamprey (n =5) after slicing were transferred in a cooled recording chamber and allowed to recover at 5°C for one hour before recording. Patch electrode was prepared from borosilicate glass microcapillaries (Hilgenberg GmbH) using a two-stage puller (PP830, Narishige, Japan). Patch electrodes (8–12 MΩ) were filled with an intracellular solution of the following composition (in mM): 130 K-gluconate, 5 KCl, 10 HEPES, 4 Mg-ATP, 0.3 Na-GTP, and 10 phosphocreatine sodium salt. The pH of the solution was adjusted to 7.4 with KOH and osmolarity to 270 mOsml^-1^ with water. Cells ventral to the central canal were recorded in whole-cell in current-clamp mode using a Multiclamp 700B amplifier (Molecular Devices Corp., CA, USA). Bridge balance and pipette capacitance compensation were adjusted and signals were digitized and recorded using Clampex software and analysed in Clampfit (pCLAMP 10, Molecular Devices, CA, USA). The neurons were visualized with DIC/infrared optics (Olympus BX51WI, Tokyo, Japan). Resting membrane potentials were determined in current-clamp mode during whole-cell recording. The following drugs were added to the extracellular solution and applied by bath perfusion: the GABAA receptor antagonist gabazine (20 mM, Tocris, Ellisville, MO, USA), the glutamate receptor antagonist kynurenic acid (2 mM, Tocris, Ellisville, MO, USA). Neurons were intracellularly labelled by injection of 0.5% Neurobiotin (Vector Laboratories) during whole-cell recordings. After recording the spinal cord slices were fixed with 4% formalin. To investigate whether the intracellularly Neurobiotin-labeled CSF-c cells express TH, the slices were incubated overnight with a mouse monoclonal anti-TH antibody and then rinsed thoroughly in 0.01 M PBS and incubated with a mixture of Alexa Fluor 488-conjugated streptavidin (1:1000, Jackson ImmunoResearch) and Cy3-conjugated donkey anti-mouse IgG (1:500, Jackson ImmunoResearch) for 3 hours at room temperature.

### *In situ* hybridisation

Lamprey (n=2) were deeply anesthetised as described above and the rostral spinal cord was removed, fixed in 4% formalin in 0.1 M PB overnight at 4°C, and then cryoprotected in 20% sucrose in 0.1 M PB. Then, 10–20 μm thick cryostat sections were made and stored at −80°C until processed. Single-stranded digoxigenin-labelled sense and antisense pkd2l1 riboprobes were generated by *in vitro* transcription of the previously cloned pkd2l1 cDNA using the Digoxigenin RNA Labeling kit [9] (catalog #11 277 073 910; Roche Diagnostics). Briefly, sections were incubated for 1 hour in prehybridisation mix (50% formamide, 5×SSC, 1% Denhardt’s, 50 g/ml, salmon sperm DNA, 250 g/ml yeast RNA) at 60°C. Sections incubated with the heat-denatured digoxigenin-labelled riboprobe were hybridised overnight at 60 °C. Following the hybridisation, the sections were rinsed twice in 1×SSC, washed twice in 1×SSC (30 min each) at 60°C and twice in 0.2×SSC at room temperature. After blocking in 0.5% blocking reagent (PerkinElmer), the sections were incubated overnight in anti-DIG antibody coupled to HRP (1:2000; RRID: AB_514497; Roche Diagnostics) at 4°C. The probe was then visualised by TSA Cy3 Plus Evaluation Kit (catalog #NEL763E001; PerkinElmer). The specificity of the hybridisation procedure was verified by incubating sections with the sense riboprobe (data not shown). The sections were rinsed thoroughly in 0.01 M PBS and then incubated with a mouse monoclonal anti-TH antibody (1:200; Millipore) overnight at 4°C, rinsed in PBS, and incubated with Alexa Fluor 488-conjugated donkey anti-mouse IgG (1:200; Jackson ImmunoResearch) and 640/660 deep-red fluorescent Nissl stain (1:1000; Invitrogen) for 2 hours and mounted with Mowiol. All primary and secondary antibodies were diluted in 1% BSA and 0.3% Triton X-100 in 0.1 M PB.

## Acknowledgements

We thank the ALM facility at the Science for Life Laboratory for access to the STED Leica Sp8.

We thank the the SciLifeLab SFO funding, the GGS foundation and the Swedish Foundation for Strategic Research (project FFL15-0031) for supporting the research.

## Author Contributions

I.T. designed and supervised the research. E.J. performed the experiments and analysed the data. J.A. performed the STED experiments and data analysis. G.C. provided guidance on sample preparation and image analysis. S.E. performed the light sheet recordings and data analysis. B.R. and S.G. provided guidance on the spinal cord neurobiology and data interpretation. I.T., E.J. and S.G. wrote the paper with contribution from all the authors.

## Data availability

All data will be available upon request.

