## supplementary_text for "ExSTED microscopy reveals contrasting functions of dopamine and somatostatin CSF-c neurons along the central canal"

**Table of contents**

**Extended Data Fig. 1. Dopaminergic CSF-c neurons are sensitive to fluid movement.**

**Extended Data Fig. 2. 3DExM of expanded cilia protruding to the central canal.**

**Extended Data Fig. 3. PKD2L1, ASIC3, and ARL13b expression on CSF-c neurons on mouse spinal cord.**

**Supplementary Movie 1. 3D ExLSM explore spatial organization of somatostatin and dopaminergic CSF-c neurons in the spinal cord.**

**Supplementary Movie 2. 3D ExSTED visualize CSF-c neurons and their cilia within the 3D geometry of the central canal.**

**Supplementary Movie 3. CSF-c neurons might contain two cilia.**

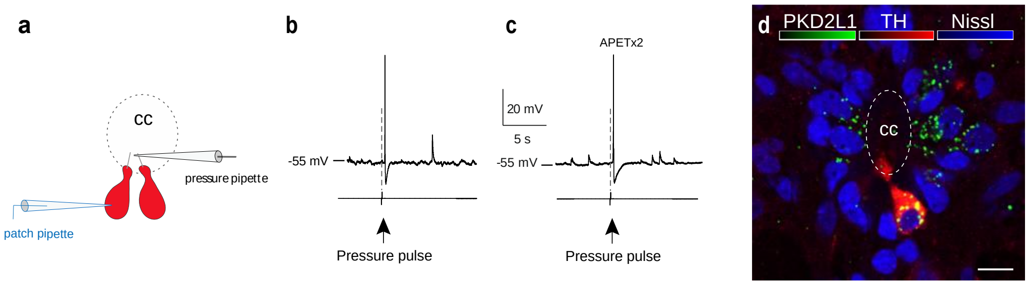


**Extended Data Fig. 1. Dopaminergic CSF-c neurons are sensitive to fluid movement**. **a**, A schematic illustration showing a dopaminergic CSF-c neuron (red) was patched and a pressure pipette was placed close to its bulb-like ending **b**, A short pressure pulse (20 p.s.i., 80 ms) elicited action potential responses in a dopaminergic CSF-c neuron. **c**, Elicited action potential responses in a dopaminergic CSF-c neuron by pressure pulse in present of the ASIC3 blocker APETx2 (2 µM). **d**, *In situ* hybridisation image showing ventrally located CSF-c neurons co-expressing the PKD2L1 channel (green), TH (red), and Nissl (blue). Scale bar, 10 µm. cc, central canal.

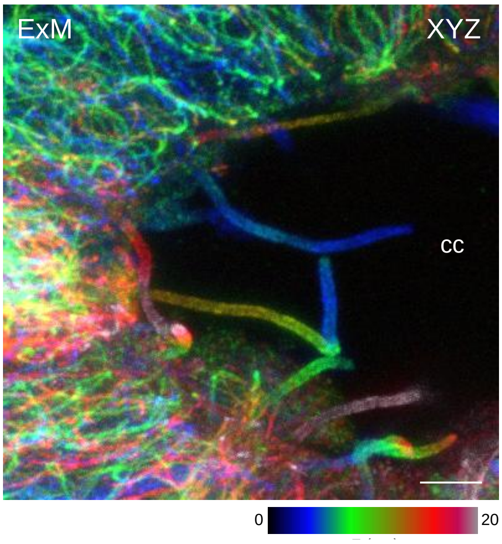


**Extended Data Fig. 2. 3DExM of expanded cilia protruding to the central canal**. Z-stack projection of expanded cilia (stained with α-tubulin) protrusion to central canal from CSF-c neurons. The z-position is indicated by the colour-coding. Scale bar, 5 µm. cc, central canal.

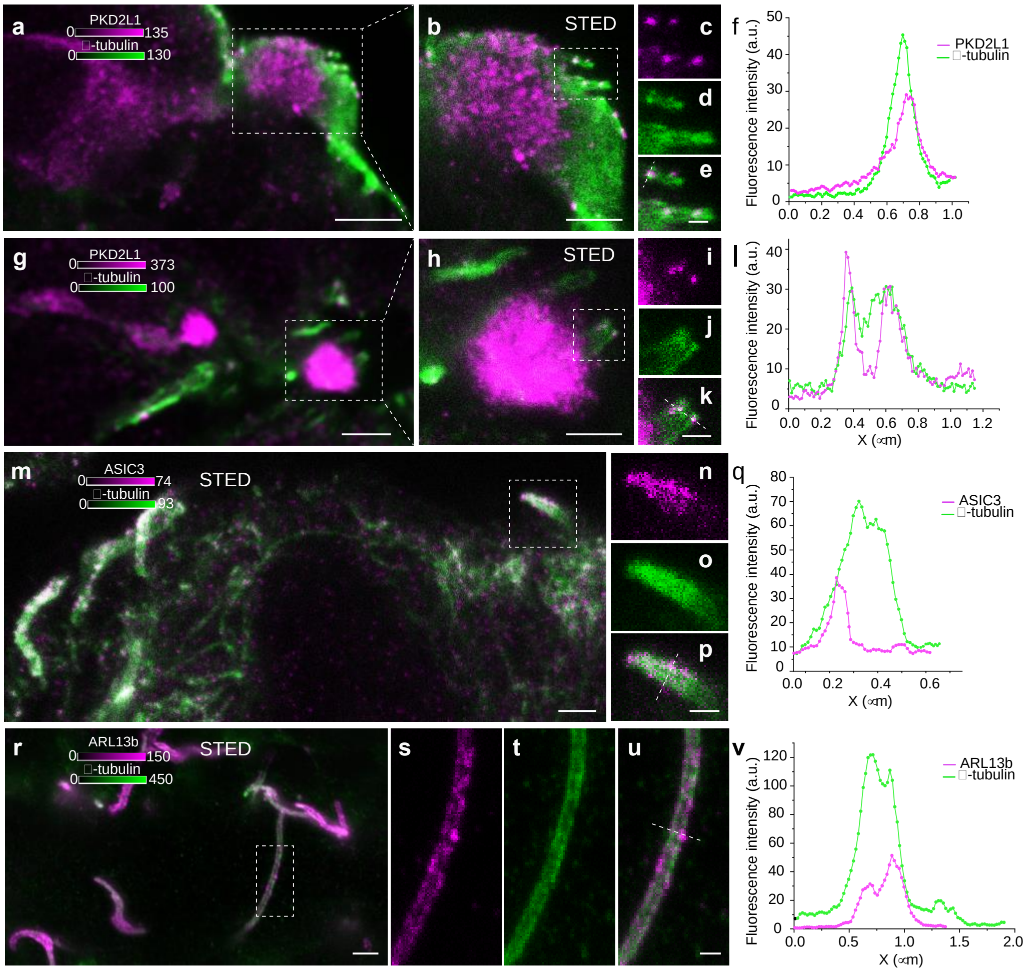


**Extended Data Fig. 3. PKD2L1, ASIC3, and ARL13b expression on CSF-c neurons on mouse spinal cord. a-l**, Two different CSF-c neurons showing immunoreactivity to the PKD2L1 channel (magenta) on their cilium stained with α-tubulin (green). Scale bar, a,g, 2 µm and b,h, 1 µm. **c-e**, Selected ROI from **b** and **i-k**, selected ROI from h is shown at higher magnification with STED microscopy with their corresponding line profiles **f** and **l**, respectively. Scale bar c-e, i-k, 300 nm. **m-p**, ASIC3 (magenta) expression on cilia of CSF-c neurons showing with STED microscopy. Scale bar, m, 1µm. **n-p**, Selected ROI from **m** is shown at higher magnification with a corresponding line profile in **q**. Scale bar, 300 nm. **r-u**, Cilia in the mouse central canal stained with α-tubulin (green) and ARL13b (magenta). Scale bar, r, 1 µm. **s-u**, STED images of a ROI from **r** showing ARLB13b expression on a cilium with corresponding line profile in **v**. Scale bar, 500 nm.

**Supplementary Movie 1. 3D ExLSM explore spatial organization of somatostatin and dopaminergic CSF-c neurons in the spinal cord.** Z-stack recorded with a light-sheet of an expanded sample of a lamprey spinal cord tissue. The reconstruction highlights specific location of somatostatin (magenta) and dopamine (green) expressing CSF-c neurons in the 3D architecture along the spinal cord.

**Supplementary Movie 2. 3D ExSTED visualize CSF-c neurons and their cilia within the 3D geometry of the central canal.** Z-stack recorded with STED of an expanded lamprey spinal cord tissue showing cilia (stained with α-tubulin) protrusion to central canal from CSF-c neurons with high resolution.

**Supplementary Movie 3. CSF-c neurons might contain two cilia**. Z-stack recorded with confocal microscopy of an expanded lamprey spinal cord tissue (3D ExM) showing of an CSF-c neuron with two cilia on its bulb. Cilia were stained with α-tubulin.
